# Extracellular lactate as an alternative energy source for retinal bipolar cells

**DOI:** 10.1101/2023.07.10.548331

**Authors:** Victor Calbiague Garcia, Yiyi Chen, Bárbara Cádiz, François Paquet-Durand, Oliver Schmachtenberg

## Abstract

Retinal bipolar and amacrine cells receive visual information from photoreceptors and participate in the first steps of image processing in the retina. Several studies have suggested the operation of aerobic glycolysis and a lactate shuttle system in the retina due to the high production of this metabolite under aerobic conditions. However, whether bipolar cells form part of this metabolic circuit remains unclear. Here, we show that the lactate transporter MCT2 is functionally expressed in inner retinal neurons and demonstrate their ability to consume extracellular lactate as an alternative to glucose. In rod bipolar cells, lactate is used to maintain homeostasis of various ions and electrical responses. Overall, our data contribute to a better understanding of inner retinal metabolism.

## Introduction

The retina is a neural tissue in which visual signals are transduced by photoreceptors and subsequently processed by inner retinal neurons. The constant turnover of the photopigment in the outer segments (OS) and the large ATP consumption required to keep the ion pumps working to maintain the dark current in the inner segment (IS) of photoreceptors are among the main reasons why the retina is the most energy-demanding neural tissue ^1–4^. Photoreceptors receive glucose, oxygen, and other metabolites from the choroidal vasculature ^5^ through the retinal pigment epithelium ((RPE; ^6^). However, the inner retina may either be vascular or avascular. In avascular retina (*e.g.,* guinea pig, rabbit), bipolar cells (BCs) and amacrine cells (ACs) probably rely only on Müller cells (MCs) as energy providers, while in the vascular retinas (*e.g.,* rodents and primates), these neurons can take up fuel directly from the inner retinal capillaries, apart from MCs and astrocytes ^5, 7^.

Interestingly, Otto Warburg’s data from 1924 suggested that tumors and the retina mostly utilize aerobic glycolysis, converting almost 70% of the glucose consumed into lactate ^8, 9^, and many later studies have supported the idea of a retinal lactate shuttle ^10–12^. Although the specific site and metabolic conditions at which lactate is released are still unclear, these investigations support the notion of a “metabolic ecosystem” between MCs, RPE, and photoreceptors, in which lactate produced by one cell type may be consumed by other cell types. However, the dynamics and extension of this shuttle to the inner retina remain a matter of debate, especially in the vascular retina ^7^.

Since the principal function of neurons is to propagate electrical signals, it is intuitive to think that their functions depend on metabolism and how these cells obtain energy ^13^. Accordingly, the observation in rats that inhibition of monocarboxylate transporters (MCTs) attenuates the ERG b- wave and oscillatory potentials ^14^. Since the ERG b-wave is generated by inner retinal neurons this data indicates the importance of lactate production for electrical signaling in the inner retina. This alteration could be partially ameliorated by exogenous lactate, revealing a potential for lactate uptake by the inner retinal neurons ^14^. This idea is supported by another study in mice, where the genetic deletion of PKM2 (aerobic glycolysis marker), which is expressed in photoreceptors, decreased the ERG a-wave and b-wave ^15^, suggesting that photoreceptor lactate is consumed in the inner retina, and is needed to maintain these ERG component.

Taking these prior data into account, we hypothesize a consumption of lactate by inner retinal neurons to maintain their physiological activity. Yet, single-cell studies to determine and support what substrate each cell type consumes under physiological conditions are still lacking.

Thus, we set out to test the role of extracellular lactate as a possible alternative energy substrate for mouse retinal bipolar cells. To this end, we expressed genetically encoded FRET nanosensors that allowed us to qualitatively determine the levels of some metabolites in real-time ^16, 17^. The putative role of lactate as a substrate for the physiological activity of inner retinal neurons was tested by calcium imaging and electrophysiological recordings. These measurements were complemented by markers of specific enzymes involved in aerobic glycolysis, pharmacological inhibition of lactate transporters and enzymes related to aerobic glycolysis.

## Results

### The inner retina expresses monocarboxylate transporters

To elucidate the potential role of extracellular lactate in the inner retina, we examined the expression patterns of MCT2 in the retina of wild-type mice. MCTs are proton-linked plasma membrane transporters that allow the transport of lactate and pyruvate into and out of cells ^18^. We focused on the MCT2 isoform because it has been reported to be the neuronal importer of lactate ^19^, especially in high-lactate environments ^20^. Immunofluorescence localization indicated that MCT2 was expressed in a subset of somas in the outer nuclear layer (ONL), likely cones, but more strongly and abundantly found in cell bodies of the inner nuclear layer (INL, Fig. 1a). Co- immunostaining with different markers indicated that MCT2 was expressed in rod bipolar cells (RBCs) and ACs in the INL (Fig. 1a).

**Fig. 1.**
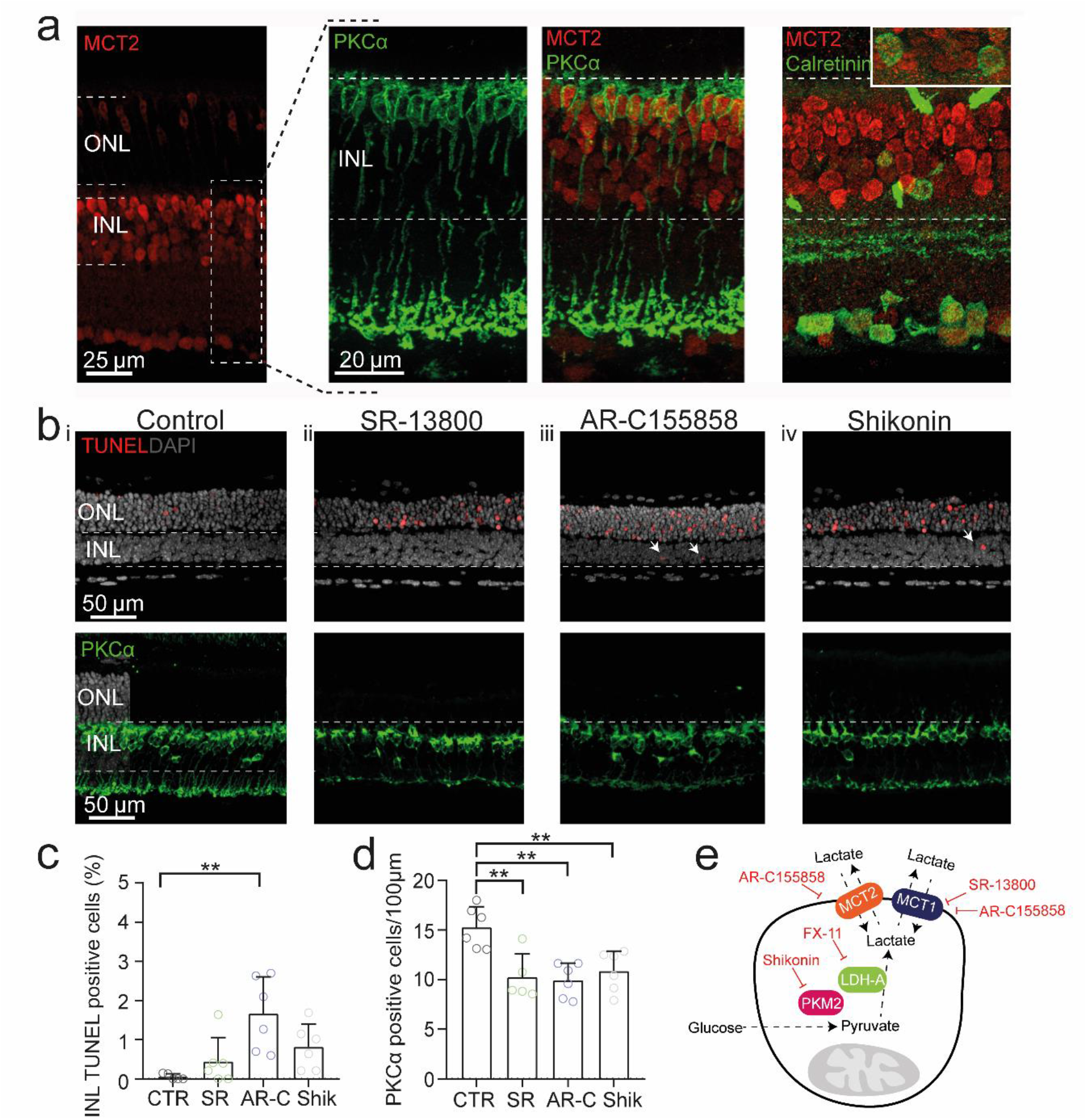
Lactate metabolism is required for inner retinal neuron survival. **a**, Immunofluorescence labeling of retinal transverse cryosections showing MCT2 expression mainly in the inner nuclear layer (INL). Co- labelling for PKCα and calretinin illustrating MCT2 expression in RBCs and a subset of ACs. **b_i_-b_iv_ (top), c**, Statistical analysis, and quantification of the cell death assay (TUNEL), performed in organotypic retinal explants, showing occasional TUNEL positive nuclei (arrowheads). When compared to control (CTR) treatment with AR-C and Shikonin increased INL cell death, while SR treatment did not. The data were analyzed with the Kruskal-Wallis and Dunn’s multiple comparison post hoc tests. **b_i_-b_iv_ (bottom), d**, Quantification of RBCs per 100 µm retinal length revealed a significant density reduction of this cell type after treatment with SR, AR-C, and Shikonin, supporting a dependence of RBCs on extracellular lactate. The data were analyzed by one-way ANOVA with Tukey’s multiple comparison post hoc test. Each dot reflects a single retinal explant. **e**, Schematic summary, showing the transporters, applied drugs used throughout the investigation, and their respective effects on lactate metabolism. Shikonin and FX-11 inhibit lactate synthesis directly, while AR-C155858 and SR-13800 block lactate transport. Graphs display mean values ± SD; asterisks indicate \**p*<.05, \*\**p*<.01. ONL = Outer nuclear layer; PKCα = Protein kinase Cα; MCT1 = Monocarboxylate transporter 1; MCT2 = Monocarboxylate transporter 2; PKM2 = Pyruvate kinase M2; LDH- A = Lactate dehydrogenase A. SR = MCT1 inhibitor; AR-C = MCT2 inhibitor; Shikonin = PKM2 inhibitor.

### Inhibition of lactate transport increases cell death in the inner retina

The widespread expression of MCT2 in the INL led us to hypothesize that extracellular lactate may be used by inner retinal cells to meet their physiological demands. To study the role of extracellular lactate in the inner retina with a functional approach, we cultured mouse organotypic retinal explants for 7 days to allow us to perform experiments under fully controlled *in vitro* conditions ^21^. We first used the TUNEL assay to examine whether MCT2 function was essential for cell survival in culture, quantifying the cell death rate as the fraction of dying cells in the INL after 4 days of treatment. Since there are no commercial inhibitors selective for only MCT2, we treated retinal explants with combinations of MCT1 or MCT1/MCT2 inhibitors, to separate the roles of MCT1 and MCT2.

In the control condition, the cell death rate was 0.06 ± 0.07% (n = 5), and a comparatively low rate (0.44 ± 0.61%, n = 6, *p* = 0.6347) was found in the INL when we blocked MCT1 with the potent and selective drug SR-13800 (SR, Fig. 1b, c) ^22^. On the other hand, when we used AR-155858 (AR-C), a potent inhibitor of both MCT1 and MCT2 ^23^, a significant increase (1.67 ± 0.94%, n = 6, *p* = 0.0014) of TUNEL-positive cells was observed (Fig. 1b, c), indicating that MCT2 function was relevant for the survival rate of INL cells.

To test the consequences of a reduction of retinal lactate production, we inhibited the enzyme pyruvate kinase (PK) M2 with Shikonin ^24^. Pyruvate kinase is a key glycolytic enzyme that controls the final step of glycolysis, converting phosphoenolpyruvate to pyruvate, and the M2 isoform is a key regulator of aerobic glycolysis specifically associated with tumor cells ^25^. The treatment with Shikonin resulted in a small increment (0.82 ± 0.57%, n = 6, *p* = 0.0502) (Fig. 1b, c). Notably, under all conditions, the number of TUNEL-positive cells in the ONL was much higher: SR = 19.9 ± 6.9%, n = 6, *p* < 0.0001, AR-C = 16.6 ± 5.2%, n = 6, *p* = 0.0007, Shikonin = 14.5 ± 5.9%, n = 6, *p* = 0.0018, reflecting the sensitivity of photoreceptors to metabolic disruptions ^6^.

After confirmation of MCT2 expression in RBCs, we set out to explore the effects of the inhibition of lactate transport and production on the survival rate of this specific cell type. To this end, the number of RBCs per 100 µm retinal section length was counted, yielding 15.3 ± 2.1 cells (n= 5, Fig. 1b, d) in untreated control explants. Interestingly, in all treatments, the RBC number decreased compared to the control condition: SR (10.2 ± 2.3, n = 6, *p* = 0.0034), AR-C (9.9 ± 1.7, n = 6, *p* = 0.0012), and Shikonin (10.8 ± 2.0, n = 6, *p* = 0.0070), suggesting that lactate production and transport were important for maintaining RBCs alive.

Taken together, these results indicated that inner retinal cell survival is sensitive to the inhibition of lactate metabolism, and specifically, that RBCs need lactate for survival in culture.

### Lactate transport is important to maintain neuronal depolarization in the inner retina

Given that the principal function of neurons is propagating electrical signals, it can be assumed that this activity depends to some degree on their energy metabolism. Yet, whether inner retinal cells use lactate apart from glucose to fuel their physiological activity remains unclear. To address this question, we performed calcium imaging experiments, which allows for a global assessment of retinal function. Retinal slices from p30 mice were incubated for 15 min with three MCT inhibitors (Fig. 2a, b), and the change in the relative fluorescence intensities of BC and AC cell bodies in the INL was measured as the amplitude and area under the fluorescence intensity curve (AUC) of the responses after a slight depolarization through 12 mM K^+^ stimulation (Fig. 2b). Under blockage of MCT1 (n = 3 retinas, ROIs = 24) and MCT1/MCT2 (n = 3 retinas, ROIs = 30), the calcium signal amplitudes were reduced to 44.5 ± 19.0% (*p* = 0.0028) and 58.5 ± 45.0% (*p* = 0.0004) of controls (Fig. 2b, c), respectively. Interestingly, acute inhibition of PKM2 with Shikonin (n = 3, ROIs = 20) had no noticeable impact on the amplitude of the calcium responses (Fig. 2b, c, 84.1 % ± 32.4, *p* = 0.9987), suggesting that Shikonin application may become effective only after prolonged treatment (*e.g.,* incubation of retinal explants).

**Fig. 2.**
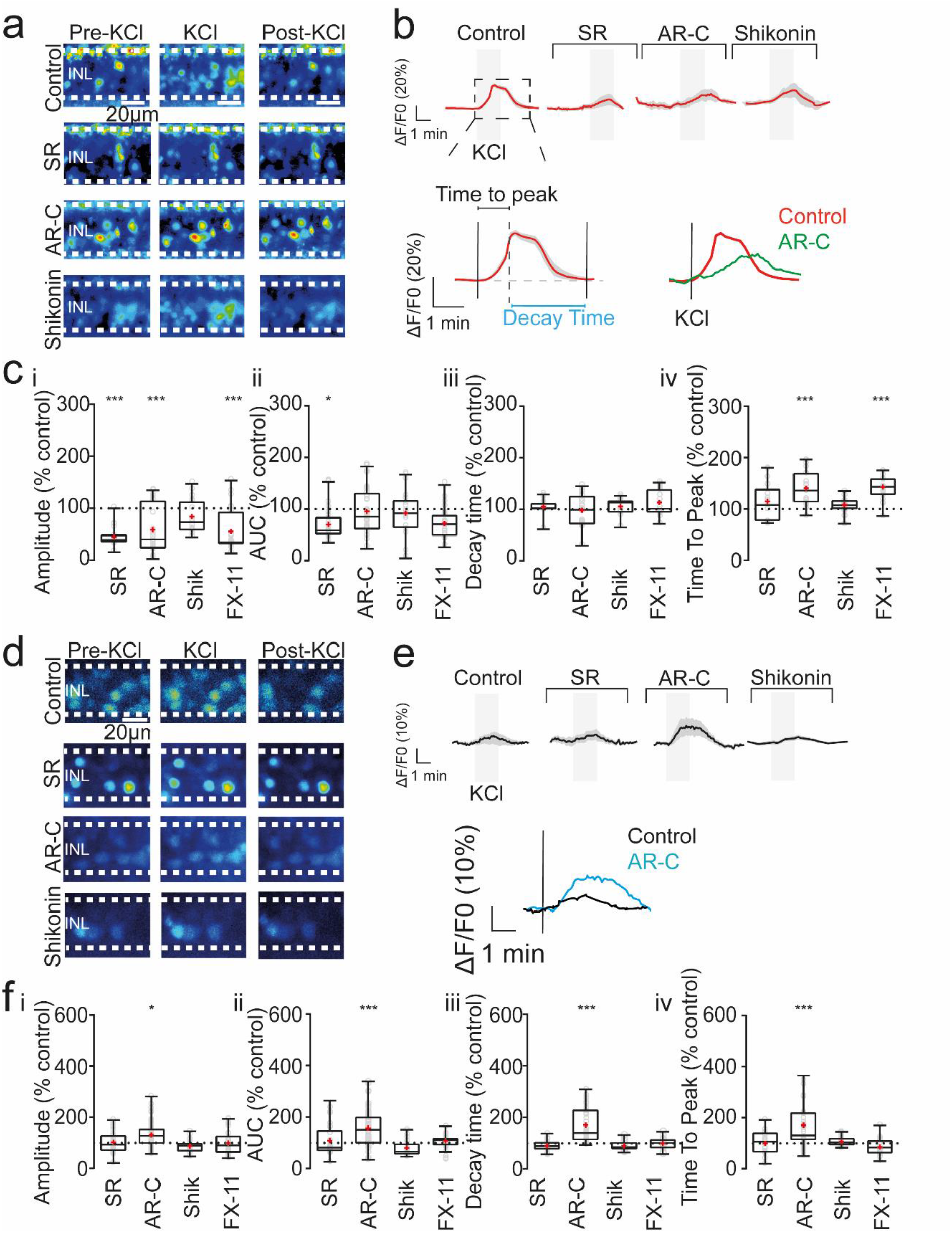
Depolarization of inner retinal cells is reduced and slowed by inhibition of lactate metabolism. **a, d**; Representative images of Fluo4-AM and Corona Green-loaded cells in the INL, before and at two time points after bath perfusion with KCl, in controls and retinas incubated with SR, AR-C and Shikonin. **b, e** (top); Traces of individual experiments in each condition (red line = mean, grey shadow = SD). **b, e** (bottom); overview of the kinetic parameters measured in the imaging responses. **c_i-iv_**; Statistical analysis of the different parameters in the calcium imaging experiments. **f_i-iv_**; Statistical analysis of the sodium imaging experiments. Box plots display the median ± min and max values and the mean in red. Individual values are displayed as open circles (gray). The control is represented as a dashed line in 100%, and results are showed as percentage of control. Asterisks indicate * *p* < .05, \*\**p* < .01, \*\*\**p*< .001. INL= Inner nuclear layer; SR= MCT1 inhibitor; AR-C= MCT2 inhibitor; Shikonin= PKM2 inhibitor; FX-11= LDH-A inhibitor. See also Figures S1 and S2.

Subsequently, we set out to test the role of lactate production in the retina through inhibition of the key enzyme lactate dehydrogenase A (LDH-A) with the drug FX-11. This enzyme is directly responsible for the conversion of pyruvate to lactate. Just as PKM2, it is a fundamental regulator of aerobic glycolysis and is frequently overexpressed in tumor tissue ^26^. Inhibition of LDH-A should decrease the aerobic glycolysis rate, and thereby lactate levels. Under this condition (n = 3, ROIs = 23), the response amplitude was again reduced compared to controls (Fig. 2c; 55.1 ± 39.9%, *p* = 0.0001). In contrast, the AUC did not change in the conditions of MCT blockage with either AR- C (Fig. 2c, 95.6 ± 43.1%, *p* = 0.9954), Shikonin (Fig. 2c, 92.2 ± 43.6%, *p* = 0.9675), and FX-11 (Fig. 2c, 71.9 ± 30.4%, *p* = 0.2046), and was only affected in the SR condition (Fig. 2c, 69.9 ± 30.2%, *p* = 0.0148), suggesting that the responses kinetics were differentially affected by the drugs. To further analyze this hypothesis, we measured the time-to-peak and the decay time of the responses. We observed an increase in the time-to-peak when MCT1/MCT2 were inhibited (Fig. 2b, c, 140.9 ± 31.7%, *p* < 0.0001) and FX-11 (Fig. 2c, 143.2 ± 23.8%, *p* < 0.0001), while MCT1 inhibition (114.9 ± 34.5%, *p* = 0.2514) and PKM2 (108.0 ± 15.8%, *p* = 0.5138) had no effect. However, the decay time was unaffected in all conditions (Fig. 4c): SR (105.0 ± 13.8%, *p* = 0.7238), AR-C (98.2 ± 30.8%, *p* = 0.9954), Shikonin (105.6 ± 15.6%, *p* = 0.9991), and FX-11 (113.4 ± 25.5%, *p* = 0.7323). Altogether, these results indicate that INL cells are susceptible to inhibition of lactate synthesis and transport, causing a disruption of their calcium responses to transient depolarization.

The unaffected response decay time was initially surprising because a disruption of energy consumption should affect the restoration of ion gradients after depolarization ^27^. However, because the calcium concentration is critical for cells, they have redundant intracellular buffers, which can quickly restore intracellular calcium levels ^28^. Therefore, the next step was to directly evaluate the ion pump function. We used the sodium probe Corona Green (Fig. 2d, e), to assay the importance of lactate on Na^+^ flux. Yet, neither inhibition of MCT1 (n = 3, ROIs = 33, 102.1 ± 40.9%, *p* >0.9999), PKM2 (n = 3, ROIs = 20, 87.6 ± 26.1%, *p* >0.9999), nor inhibition of LDH-A (n = 3, ROIs = 29, 100.0 ± 45.7%, *p* > 0.9999) caused significant effects (Fig. 2f). Similar results were observed for the AUC: SR (108.9 ± 64.2%, *p* > 0.9999), Shikonin (80.7 ± 30.9%, *p* = 0.3175), and FX-11 (108.3 ± 23.1%, *p* = 0.9649). However, when we inhibited MCT1/MCT2 with AR-C, the amplitude (131.8 ± 51.5%, *p* = 0.0071) and AUC (157.1 ± 70.9%, *p* = 0.0002) of the sodium responses increased significantly (Fig. 2f).

Similar results were obtained regarding the other response kinetic parameters: neither MCT1 (100.3 ± 41.7%, *p* > 0.9999), PKM2 (107.2 ± 19.4%, *p* > 0.9999), nor LDH-A (85.9 ± 31.7%, *p* = 0.4242) inhibition caused changes in the time-to-peak. When we analyzed the decay time, no changes were observed either: MCT1 (90.9 ± 21.5%, *p* = 0.8233), PKM2 (89.7 ± 18.9%, *p* = 0.6635), and LDH-A (99.9 ± 26.3%, *p* > 0.9999). But when we inhibited both MCT1/2 with AR-C significantly affected the time-to-peak (171.0 ± 85.4%, *p* < 0.0001), and the response decay time increased (171.0 ± 70.2%, *p* < 0.0001) (Fig. 2f).

Taken together, these results indicate that MCT2 inhibition was necessary to affect the response kinetics, suggesting an alteration in ion pumping to restore sodium homeostasis ^27, 29^. If this idea was indeed correct, changes in the basal levels of calcium and sodium should also be observable. Therefore, we measured basal fluorescence levels and calculated the slope of the relative fluorescence to analyze whether calcium or sodium accumulation occurred under inhibition of the different transporters and enzymes related to aerobic glycolysis.

As for the relative calcium levels, it was striking to see that under three different conditions (*i.e.,* SR, Shikonin, and FX-11), the relative levels decreased significantly: Blockage of MCT1 caused a reduction of 74.8 ± 43.9% (*p* < 0.0001), whereas inhibition of PKM2 and LDH-A produced a reduction of 77.1 ± 85.6% (*p* < 0.0001) and 147.1 ± 78.0% (*p* < 0.0001), respectively (Fig. S1). In contrast, incubation with AR-C resulted in a slight increase in relative intracellular calcium levels (34.9 ± 79.2%, *p* = 0.0148; Fig. S1b, e).

Conversely, for relative sodium levels, only inhibition of MCT2 and, to a lesser extent, of LDH-A were effective (Fig. S2). A significant drop in the relative sodium level of 282.2 ± 155.1% (*p* < 0.0001) was seen under AR-C incubation, while FX-11 caused a minor reduction of 31.9 ± 55.7% (*p* = 0.0014) (Fig. S2). In summary, these results demonstrate that inhibition of lactate metabolism can disrupt several features of neuronal depolarization in inner retinal cells.

### Lactate can be used as an alternative fuel for RBCs

To collect further evidence for the involvement of lactate in inner retinal neuron physiology, we set out to measure representative voltage-gated currents in retinal cells performing whole-cell patch- clamp recordings under different lactate and glucose concentrations. In mammals, the concentration of lactate in the retina has been reported to fluctuate between 5 and 50 mM, depending on the species ^30^. Here, we measured the lactate released by retinal explants after 4 days in culture and obtained a concentration of 16.7 mM ± 5.7 lactate in the culture medium, in line with previously reported values ^30^.

Then, *ex vivo* retinas from p30 mice were incubated for 15 min in 4 different conditions: 20 mM glucose (control), 20 mM lactate + 10 mM mannitol (no glucose, mannitol was used for osmolarity compensation), 40 mM lactate (no glucose) and 20 mM mannitol (no glucose). Subsequently, the outward currents (*I_out_*), the calcium current (*I_Ca_*) and the membrane potential (*V_memb_*) were measured under photopic conditions.

These electrophysiological experiments showed that in retinas incubated with 20 or 40 mM lactate, RBCs did not change the amplitudes of their *I_Ca_*, *I_out_*, and *V_memb_* compared to retinas incubated with glucose (Fig. S3 and Table 1). Retinal cells incubated with 20 mM mannitol showed a decrease in *I_Ca_* and *I_out_* and an increase in *V_memb_* (Fig. S3 and Table 1). A specific feature of the calcium current in RBCs is the presence of transient reciprocal feedback from A17 cells ^31^. This feedback was only affected in the presence of 40 mM lactate and was abolished under mannitol conditions (Fig. S3 and Table 1). In contrast, the 20 mM condition maintained the amplitude of this reciprocal feedback current. These results indicate that 20 mM lactate can substitute for glucose, suggesting that RBCs can use this metabolite as an alternative substrate to maintain their physiological activity. Moreover, a saturated lactate environment and glucose deprivation conditions affect A17 feedback.

**Table 1.** Summary of the principal electrophysiological characteristics of RBCs under different metabolic conditions. Values are given as mean ± SD, and total number of cells (n).

| | $I_{out}$ (pA) | $I_{Ca}$ (pA) | Reciprocal feedback (pA) | $V_{memb}$ (mV) |
| --- | --- | --- | --- | --- |
| Glucose | 775.9 ± 73.2 (12) | 20.8 ± 3.7 (7) | 26.5 ± 10.4 (4) | -41.2 ± 5.4 (10) |
| Lactate 20mM | 729.2 ± 140.3 (9) | 19.0 ± 2.8 (8) | 23.1 ± 4.5 (6) | -41.1 ± 7.6 (9) |
| Lactate 40mM | 841.7 ± 96.7 (8) | 17.2 ± 2.1 (6) | 3.6 ± 1.7 (4) | -40.8 ± 6.5 (7) |
| Mannitol | 364.6 ± 86.7 (9) | 8.2 ± 4.3 (4) | --- (4) | -31.5 ± 6.5 (7) |
| Gluc + SR | 667.8 ± 147.9 (8) | 18.9 ± 5.5 (3) | 28.3 ± 9.9 (3) | -36.1 ± 7.4 (8) |
| Gluc + AR-C | 677.5 ± 84.3 (8) | 21.8 ± 6.7 (3) | 28.5 ± 2.1 (3) | -37.6 ± 5.3 (9) |
| Lact 20mM + SR | 633.8 ± 173.3 (8) | 20.6 ± 6.7 (5) | 28.5 ± 5.4 (5) | -35.8 ± 5.7 (8) |
| Lact 20mM + AR-C | 438.0 ± 95.9 (5) | 8.4 ± 3.8 (8) | 11.3 ± 4.8 (5) | -31.2 ± 4.9 (5) |

To test functional MCT2 expression in RBCs, retinal slices were incubated with different MCT inhibitors in the 20 mM lactate condition, because this concentration triggers an influx of this metabolite (see below). When we measured the above parameters, we observed that alterations occurred only when MCT1 and MCT2 were inhibited simultaneously, but not when only MCT1 was inhibited (Fig. 3 and Table 1). These results are in line with the immunolabeling and imaging experiments, supporting MCT2 expression in RBCs and its relevance for neuronal activity.

**Fig. 3.**
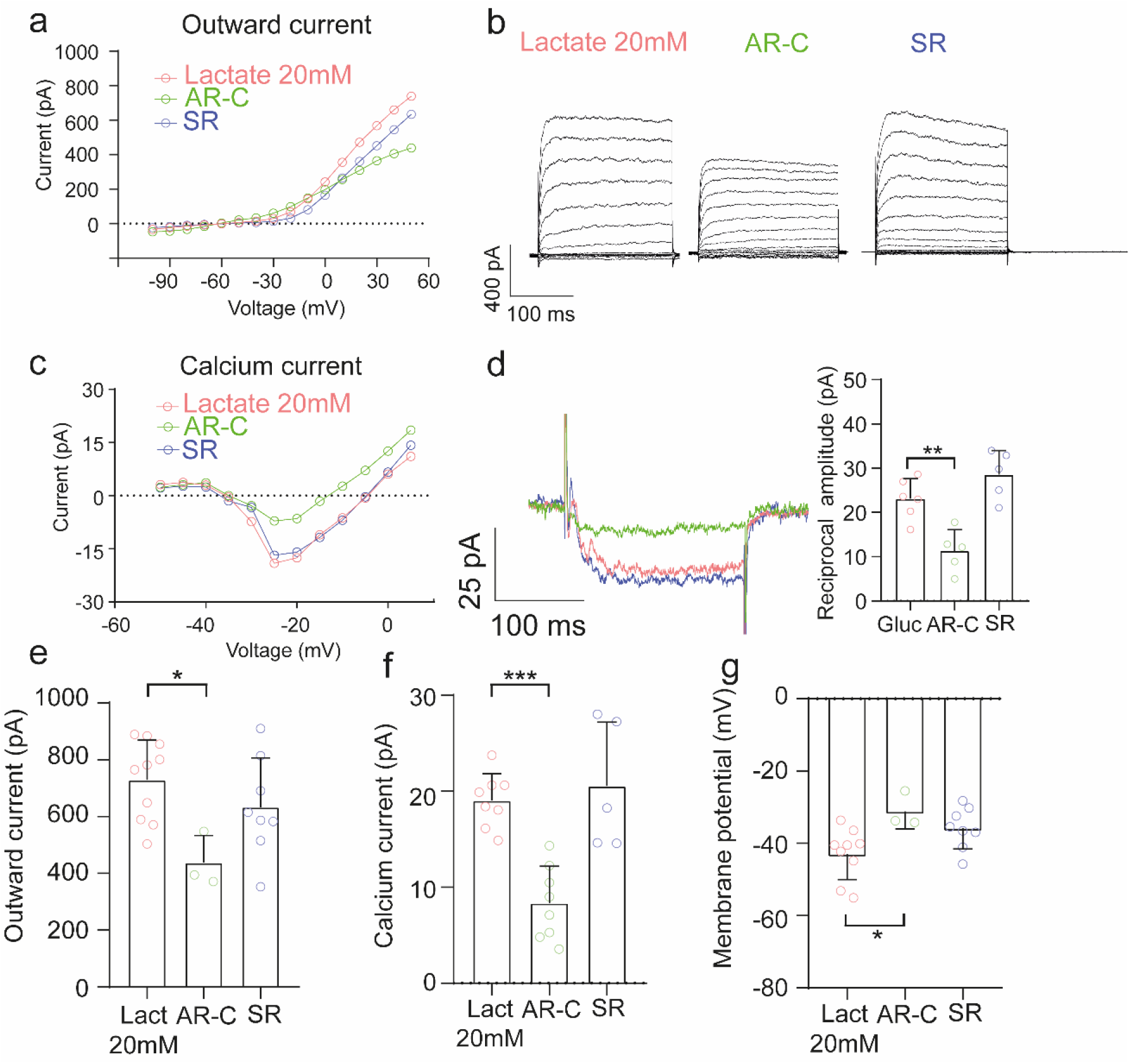
Inhibition of lactate transport through MCT2 induces alterations in RBCs currents. (a, c), Comparison of the voltage-current relationship of the outward current and calcium current under different conditions. (b, d), Representative recordings to depolarizing voltage steps. Reciprocal feedback was altered only in the AR-C condition (p = 0.0042) but was unaffected in the SR condition (p = 0.1957) compared to controls. (e, f), In the presence of 20 mM lactate, we observed a decrease in the outward current (p = 0.0224) and calcium current (p = 0.0003) only under MCT2 inhibition, but no differences were noted either in the outward current (p = 0.3925) or calcium current (p = 0.8146) when only MCT1 was blocked. (g), Similar results were obtained when we measured the membrane potential, where AR-C (p = 0.0321) caused a significant depolarization, while the SR condition was not different from controls (p = 0.0822). Data were analyzed using ordinary one-way ANOVA, followed by Tukey’s multiple comparison post-hoc test. Each dot represents a single recorded cell. Graphs represent the mean ± SD; asterisks indicate * *p* < .05, \*\**p* < .01, \*\*\**p*< .001. SR = MCT1 inhibitor; AR-C= MCT1 and MCT2 inhibitor. See also Figures S3 and S3, and Table 1.

Finally, we wondered how lactate compared to glucose as a fuel suitable to meet the physiological demands of RBCs. To this end, we applied 20 mM glucose and blocked the different MCTs. As expected, neither AR-C nor SR altered or decreased the amplitude of *I_out_*, *I_Ca_*, *V_memb_*, and reciprocal feedback currents (Fig. S3 and Table 1), suggesting that RBCs can use both glucose and lactate as alternative and equivalent fuels.

### Intracellular lactate dynamics in the inner retinal cells

Previous results have supported the idea of lactate consumption in inner retinal cells^14^. Nonetheless, to date, there is a lack of information regarding the intracellular lactate flux in the retina. For the purpose of investigating lactate dynamics in INL cells, we expressed the FRET nanosensor h-Syn-Laconic (neurons) and GFAP-Laconic (MCs) in organotypic retinal explants, as previously described ^16, 32^. First, we evaluated whether nanosensor expression was functional in inner retinal neurons, by measuring the ratio between the emissions obtained from monomeric teal fluorescent protein (mTFP)/Venus in the soma of putative BCs and ACs in the INL (Fig. S5). The lactate nanosensor was then saturated with 10 mM lactate (Fig. S4). To deplete the intracellular levels of these metabolites, we used 10 mM pyruvate to exploit the trans-acceleration property of MCTs ^33^. The delta ratio between depletion and saturation for Laconic was 20.3% ± 4.1 (Fig. S4), which is in line with the responses of the retina and other cells and tissues ^16, 32^.

Together, these results showed that FRET nanosensors can be functionally expressed in retinal explant cultures to study metabolic dynamics on a single-cell level in neurons.

After confirming the functional expression of the Laconic lactate sensor in inner retinal neurons, we aimed to verify the function of MCT2 in the INL. Since these transporters work bidirectionally, we applied 10 mM lactate stimulation to trigger lactate influx into cells, which triggered a fluorescence increase of 7.9 ± 3.4% (Fig. 4). Under inhibition of MCT1 (SR = 9.9 ± 3.6%, *p* = 0.3768) or MCT1 and MCT4 (Syrosingopine = 9.0% ± 3.1, *p* = 0.8192), this amplitude was unaffected (Fig. 4). However, when MCT1 and MCT2 were blocked, lactate influx was abolished (Fig. 4b, AR-C = -1.9 ± 0.6%, *p* = 0.0005), confirming the predominant role of MCT2 in lactate flux in inner retinal neurons. Overall, these results demonstrate the functional expression of MCT2 in BCs and ACs.

**Fig. 4.**
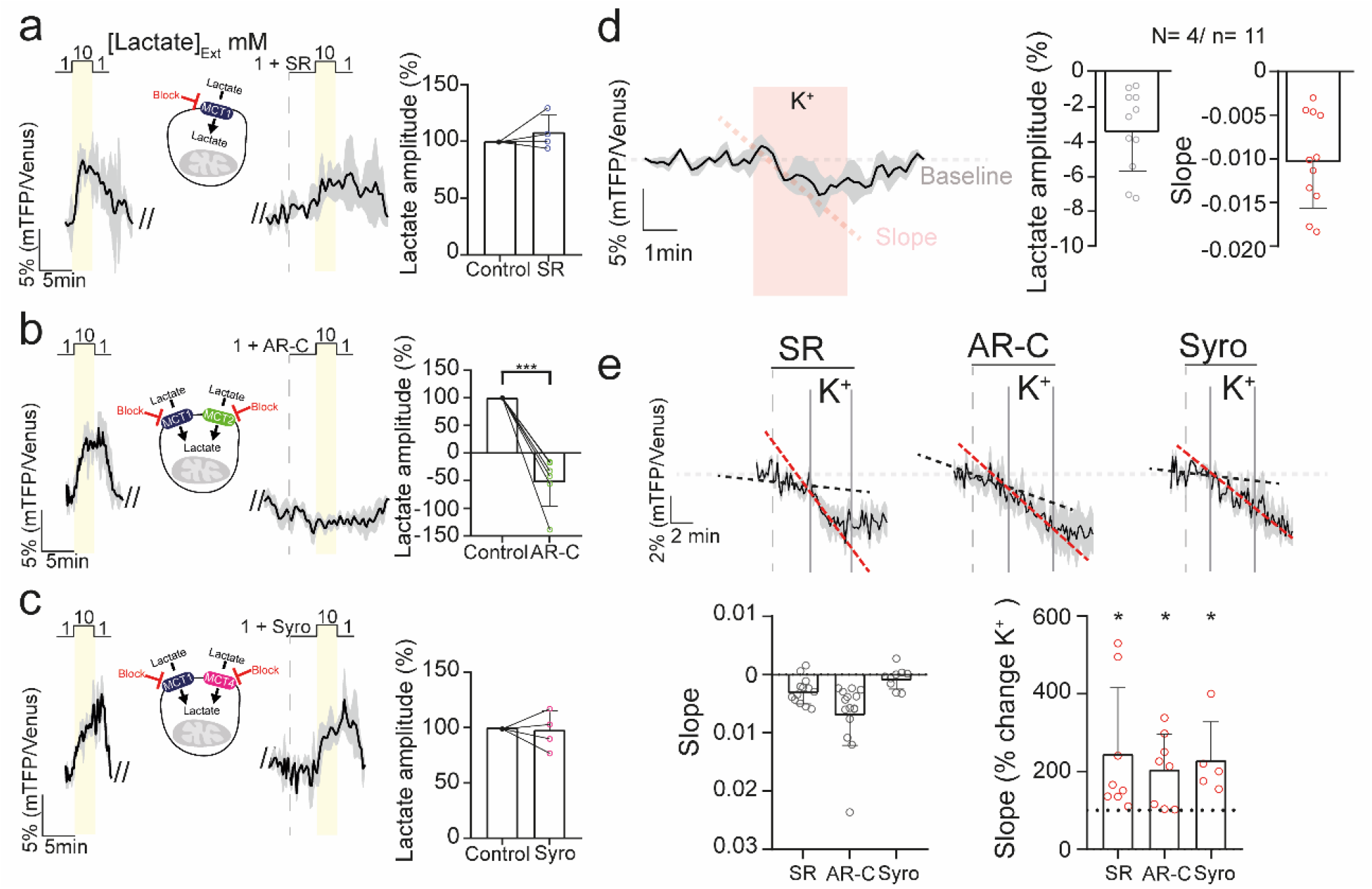
Combined MCT1/MCT2 inhibition decreases intracellular lactate levels in inner retinal neurons. **(a-c, left)** Lactate influx evoked by stimulation with 10 mM lactate. **(a-c, center)** Lactate influx responses after incubation with different drugs targeting MCTs. **(a-c, right)** Statistical analysis of the response amplitude. An evident response disruption was noted only when MCT2 was inhibited. However, under MCT1 and MCT4 inhibition, the amplitude was unaltered. **(d)** Modulation of intracellular lactate levels by transient depolarization. **(e, top)** Effects of inhibition of different MCTs and retinal depolarization on lactate dynamics. Black dashed lines indicate drug-induced change in slope and red dashed lines indicate the effect of drug + KCl. **(e, bottom)** Statistical analysis of the slope of the responses under basal conditions (only drug, left) and depolarization (drug + K+, right). The data revealed that MCT2 inhibition led to a reduction in intracellular lactate levels, resulting in lactate consumption. This consumption was exacerbated after depolarization in SR, AR-C, and Syro. Data were analyzed using either a paired Student’s t-test or Wilcoxon matched-pair test. The black trace represents the average of one experiment, whereas the light grey shadow represents the standard deviation. Graphs display the mean ± SD. * Indicates p < .05. SR = MCT1 inhibitor; AR-C = MCT1 and MCT2 inhibitor; Syro = MCT1 and MCT4 inhibitor. The number of experiments is represented as N = number of explants and n = number of cells recorded. See also Figures S5 and S6.

We then investigated how lactate dynamics change under different conditions in retinal neurons. To that end, we compared the responses under basal conditions and under moderate depolarization with 12 mM KCl. A moderate depolarization triggered a transient decrease in intracellular lactate, as evidenced by a -4.6 ± 2.0% fluorescence drop in inner retinal neurons, suggesting lactate consumption (Fig. 4d). However, it is important to note that one third of the cells recorded (35.3%) did not respond to this stimulation (Fig. S6).

To compare the lactate dynamics in different inner retinal cells, we performed the same experiment in Müller glial cells (MCs), to separate neuronal and non-neuronal components. In these cells, depolarization produced an increase in lactate levels (4.9% ± 4.1; Fig. 5b). We also expressed a glucose nanosensor (Δ6) to analyze the glucose levels in MCs. Remarkably, depolarization triggered a transient decrease in MCs intracellular glucose, as evidenced by a fluorescence drop measured using the Δ6 nanosensor (-7.5 ± 4.9%; Fig. 5a), suggesting higher glucose consumption triggered by K^+^ in MCs. This result is in line with data obtained from astrocytes in the brain ^34^.

**Fig 5.**
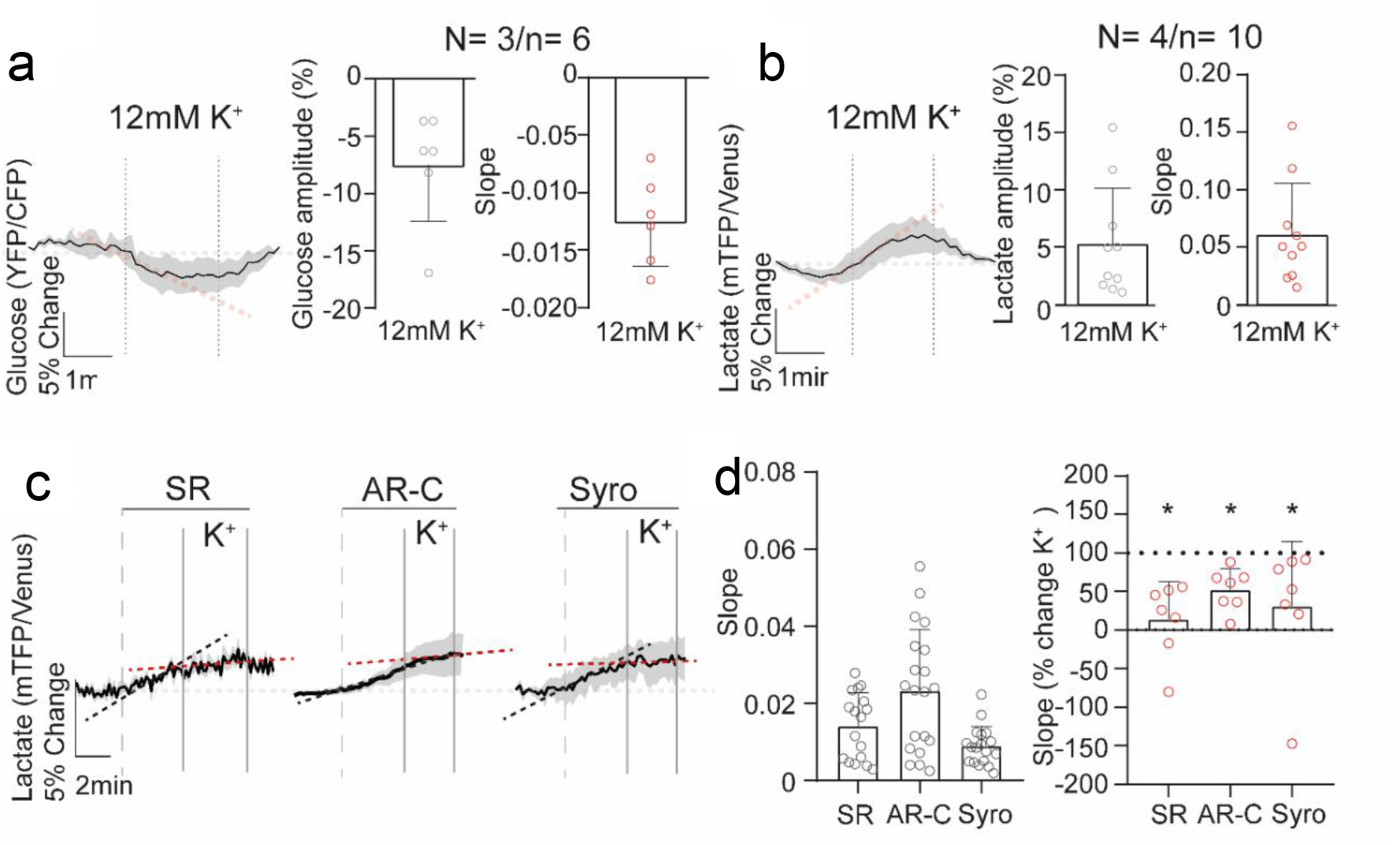
Lactate dynamics of MCs under different conditions. **(a and b)** Effect of transient depolarization on intracellular glucose (left) and lactate (right) levels. **(c)** Lactate dynamics after inhibition of different MCTs and retinal depolarization. Black dashed lines indicate drug-induced changes in slope and red dashed lines indicate the effect of drugs + KCl. **(d)** Statistical analysis of the slope of the responses in the basal condition (drug only) and under depolarization (drug + K^+^). Accumulation of intracellular lactate was observed after bath application of different MCT inhibitors. However, this increase diminished after depolarization under all conditions. Data were analyzed either with paired Student’s t-test or Wilcoxon matched-pairs test. Graphs display the mean ± SD. * Indicates *p* < .05. SR = MCT1 inhibitor; AR-C = MCT1 and MCT2 inhibitor; Syro = MCT1 and MCT4 inhibitor. The black trace represents the average response, while the light grey shadow represents the standard deviation. The number of experiments is represented as: N = number of explants; n = number of cells recorded.

Overall, these results revealed a potential difference in metabolism between neurons and MCs in the retina, where neurons may consume lactate, whereas MCs may produce it. To test this hypothesis, we performed a transport-stop protocol ^16, 35, 36^. After inhibition of MCTs, the intracellular lactate levels only decreased in the AR-C condition in neurons (Fig. 4e), confirming that inner retinal neurons consume lactate through MCT2 under basal culture conditions. Conversely, the intracellular lactate levels increased under all conditions in MCs after inhibition of MCTs (Fig. 5c, d left). This result suggests that a lactate shuttle from MCs to inner retinal neurons may operate under specific conditions ^37^.

Remarkably, when the retina was depolarized, lactate decreased significantly in neurons under all conditions, even when MCT1 (145.5% ± 169.5, *p* = 0.0456) and MCT1/MCT4 (129.9% ± 98.2, *p* = 0.0416) were inhibited. The decrease after inhibition of MCT2 (105.7% ± 90.9, *p* = 0.0393) suggested that depolarization triggered an increase in lactate consumption.

In contrast, in MCs, when the retina was depolarized, lactate increase ceased, leading to a new and higher intracellular lactate baseline (Fig. 5c, d right, SR *p* = 0.0156, AR-C *p* = 0.0156, and Syro *p* = 0.0156). These results reveal flexible and dynamic regulation of glucose and lactate consumption in the inner retina, depending on the activity status of the retina.

## Discussion

Although photoreceptors are the cells with the highest energy consumption in the retina ^3, 4^, it is equally important to understand the metabolism of inner retinal cells, which face the challenge of being further away from the main choroidal blood supply.

MCT2 expression in the inner retina is not surprising because this transporter is known as the neuronal MCT. Several authors have proposed the neuronal consumption of lactate taken up through this specific transporter isoform ^19, 38^, which is in line with prior immunohistochemical studies in the retina ^39, 40^. However, a demonstration of this transporteŕs function and its role in inner retinal physiology was missing. In vascular retinas, inner retinal cells can obtain metabolites from alternative sources ^5^, and previous studies have demonstrated the ability of the neuroretina to be metabolically flexible to meet its energy demands ^6^. Although that study focused on the outer retina, some important enzymes involved in different metabolic pathways were expressed in the INL, supporting a similar metabolic flexibility for the inner retina. Here, we found that under prolonged MCT blockage, RBCs displayed multiple functional alterations, indicating the dependence of RBCs on extracellular lactate levels under prolonged treatments.

Many studies have suggested dysregulation of calcium and sodium homeostasis under metabolic disturbances ^29, 41^, because of the strong dependence of the maintenance of ionic gradients on sustained ATP production ^42^. Since the transport of ions across biological membranes against concentration gradients is energy-intensive, a disruption in metabolism will affect the restoration of ion gradients ^27^. Indeed, a delayed return to baseline was observed here in sodium signaling, supporting this idea. This sluggish return to baseline can produce an accumulation of these ions in the cytoplasm, leading to a reduction (and slower) influx of different ions, specifically in calcium signaling ^29^. An accumulation of intracellular calcium may underlie the depolarization observed after AR-C and mannitol application in RBCs, as seen in the electrophysiology experiments, and likely is the reason for increased cell death observed after prolonged exposure in explant cultures.

Regarding the substantial increase in sodium responses, the drop in the sodium baseline could explain these results. Many studies have shown that an increase in cytoplasmic calcium concentration directly leads to an increase in mitochondrial calcium levels, which regulate numerous enzymes ^43, 44^. Subsequently, to recycle calcium ions, mitochondrial Ca^2+^ efflux is mediated by an electrogenic Na^+^/Ca^2+^ exchanger (NCLX) ^45, 46^, causing a decrease in sodium levels. In addition, this reduction in cytoplasmic sodium may be enhanced following depolarization caused by high intracellular calcium levels, leading to an upregulation of ion pumps that produce a sodium efflux ^42^. A previous study proposed non-canonical modulation of ATP levels mediated by the Na^+^/K^+^ pump, where the Na^+^ flux was capable of controlling glycolysis and ATP production ^36^. Therefore, the regulation of calcium and activation of Na^+^ pumping may induce a reduction in cytoplasmic sodium levels.

When ion gradients are affected, alterations in different electrical responses should be expected. Some studies have shown that ERG waves are sensitive to glucose deprivation and various metabolic stressors ^1, 47^. Specifically, inhibition of MCTs with α-cyano-4-hydroxycinnamic acid (4- CIN) attenuates the b-wave with delayed implicit time. However, in the presence of extracellular lactate, a partial recovery was obtained ^14^. These results suggest a possible consumption of extracellular lactate by ON BCs to maintain electrical responses. Our results support this notion because the inhibition of MCT2 caused a reduction in outward and calcium currents and depolarized the membrane potential in RBCs. Additionally, the attenuation induced in the currents and membrane potential by glucose deprivation (*e.g.,* mannitol conditions) is countered in the presence of extracellular lactate ^48^. Nevertheless, here, we show that glucose is preferred over lactate to maintain its physiological activity ^49^. Consuming glucose is essential for cells to maintain various metabolic products and by-products in equilibrium that are important for cell survival ^50^. In addition, the split between lactate and glucose consumption may allow for faster adaptation to changes in neuronal energy demand ^51^, which is crucial for the retina since this tissue has a dynamic neuronal activity depending the variation of visual stimuli.

Our results demonstrate lactate consumption by inner retinal neurons, which is exacerbated under depolarization, indicating that a retinal lactate shuttle from Muller glial cells to neurons might operate under our experimental conditions ^10^. However, we cannot rule out the possibility that other cells, such as rod photoreceptors, produce lactate under certain physiological and metabolic circumstances, as it has been recently reported ^12, 52^. Interestingly, this observation cannot be extrapolated to all inner neurons, since not all neurons displayed lactate consumption after depolarization (Fig. S6), which reflects the large heterogeneity of bipolar and other cells in the INL ^53^. In addition, a lactate consumption was also seen in some MCs (Fig. 5c, d right) suggesting that some MCs can also be lactate consumers in specific conditions ^11, 12^

Given that our experiments were carried out under photopic conditions, it is possible that this metabolic model only applies under these circumstances. This makes the retina a complex but interesting tissue to study metabolism, because different cells might have different metabolism under different light levels, and the metabolites to meet their metabolic demand could shuttle between the RPE, photoreceptors, ONL, and INL. In summary, the observations reported here suggest that the inner retina participates in a retinal lactate shuttle, where MCs release lactate under basal conditions, and inner retinal neurons potentially consume a percentage of this extracellular lactate to meet their neuronal activity (Fig. 6).

**Fig. 6.**
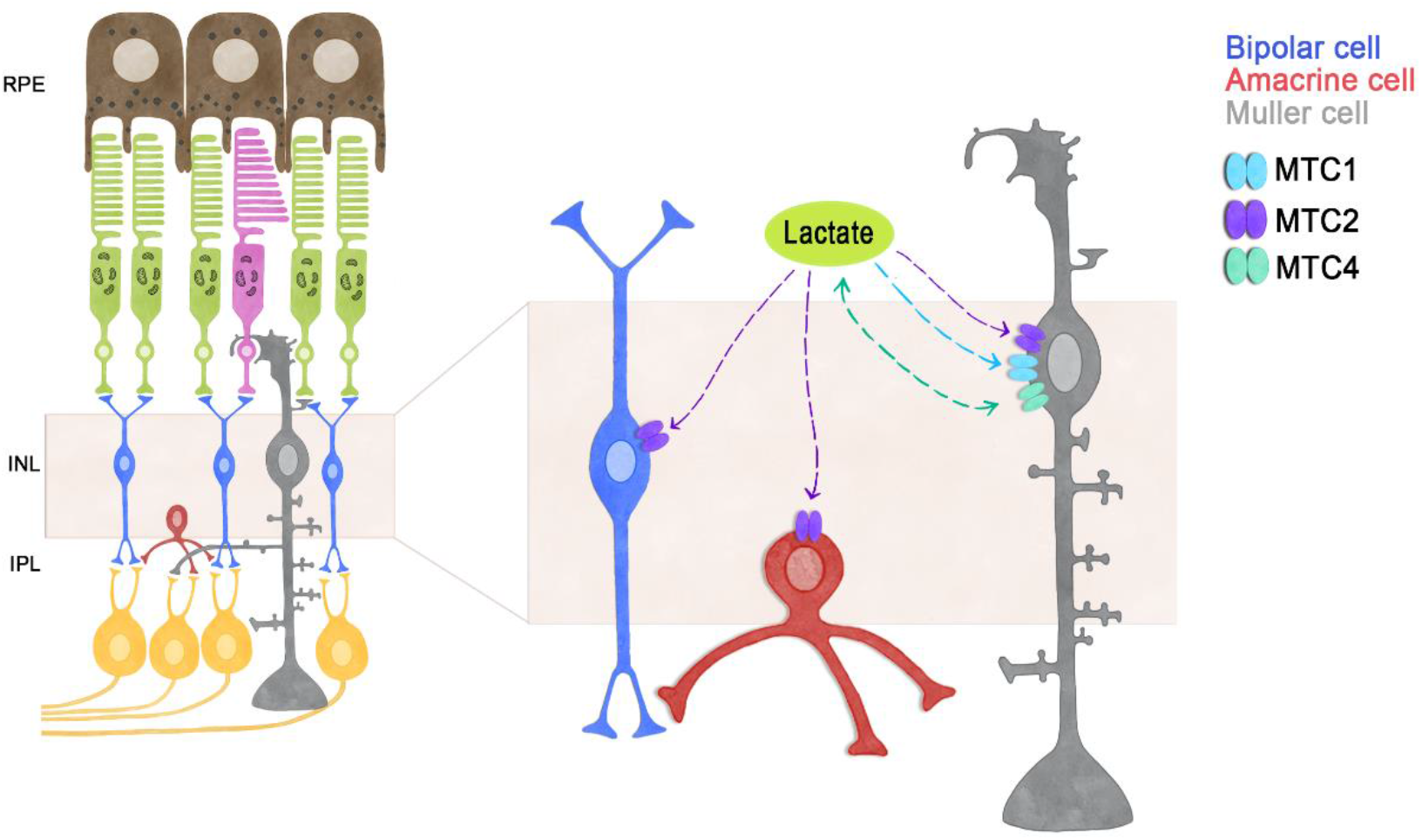
Model for the lactate dynamics in inner retinal cells. General model proposing a consumption of lactate by inner retinal cells. Previous work ^32^ demonstrated the functional expression of MCT1, MCT2, and MCT4 in MCs, where MCT2 (to a minor degree also MCT1) mainly regulates lactate influx, while MCT4 mediates lactate efflux, contributing to the accumulation of extracellular lactate. Here, we propose that this extracellular lactate produced by MCs and possibly other retinal cell types is consumed by BCs and ACs through MCT2. MCT1= Monocarboxylate transporter 1; MCT2= Monocarboxylate transporter 2; MCT4= Monocarboxylate transporter 4.

It is important to emphasize that our study has limitations. For instance, all measurements were obtained from cell bodies, whereas the long cellular projections of MCs, BCs and ACs might be metabolically isolated and could display different dynamics. Furthermore, the experimental conditions used here might not accurately reflect the physiological conditions of the *in vivo* retina. While retinas were light-adapted, photoreceptors can show variations in their activity after prolonged time in culture depending on the status of the RPE ^54, 55^.

## Author contributions

VCG and OS contributed to the conception and design of the study. VCG, YC and BC contributed to data acquisition, analysis, and figures preparation. VCG, YC, BC, FPD, and OS contributed to article writing and approved the submitted version.

## Acknowledgments

This study was supported by FONDECYT grant No. 1210790, the National Agency for Research and Development (ANID) scholarship 2018 - 21180443 (V.M.C). We wish to thank Dr. Ivan Ruminot, Dr. Felipe Baeza-Lehnert, and Dr. Felipe Barros from the Centro de Estudios Científicos (CECs) in Valdivia (Chile) for sharing their expertise in metabolic sensor FRET measurements, and Dr. Felipe Tapia from the Universidad de Valparaíso for helping with the data analysis.

## Declaration of interests

The authors declare no competing interests.

## RESOURCE AVAILABILITY

### Lead contact

Further information and requests for resources and reagents should be directed to and will be fulfilled by the lead contact, Oliver Schmachtenberg

### Materials availability

This study did not generate new unique reagents.

### Data and code availability

Data reported in this paper will be shared by the lead contact upon request. This paper does not report original code. Any additional information required to reanalyse the data reported in this paper is available from the lead contact upon request.

## EXPERIMENTAL MODEL AND STUDY PARTICIPANT DETAILS

Wild type C57Bl/6 and C3H mice were housed under standard white cyclic illumination, with water and food *ad libitum*, and were used irrespective of gender. All efforts were made to minimize the number of animals used and their suffering. Protocols compliant with the German law on animal protection and were reviewed and approved by the ‘‘Einrichtung für Tierschutz, Tierärztlicher Dienst und Labortierkunde’’ of the University of Tübingen and were approved by the bioethics committee of the University of Valparaiso, in accordance with the Chilean animal protection law No. 20.380. To isolate the eyes, animals were deeply anesthetized by inhalation using isoflurane and euthanized via decapitation.

## METHOD DETAILS

### Organotypic retinal explant culture

Retinal explants obtained from post-natal (p) day 9 wild-type mice were cultured as previously described ^56^. Briefly, the retinas were treated for 15 min with 0.12% Proteinase K (Sigma-Aldrich, Cat. No. P2308) at 37°C for isolation of the retina together with the RPE. Then, the eyes were placed for 5 min in DMEM medium with 10% fetal bovine serum (FBS) to deactivate Proteinase K. Finally, the retina was placed with the RPE facing down on cell culture inserts (Millicell, Cat. No. PICM0RG50, Merck Millipore) with DMEM culture medium (Thermo Fisher Scientific, Cat. No. 31600034), containing 10% FBS (Sigma-Aldrich) with 15 mM glucose, which was replaced every two days. The cultures were incubated at 37°C in 5% CO_2_, and 95% humidity for 14 days in a water-jacketed incubator (Thermo Scientific).

### Immunostainings

Retinas from p30 mice were fixed in 4% paraformaldehyde for 45 minutes, washed in PBS buffer, cryopreserved with solutions containing 10, 20, and 30% sucrose before embedding in tissue- freezing medium, and stored frozen at -20°C. Transverse sections of 14 µm diameter were obtained with a cryostat (Leica CM-1900) and deposited on poly-L-lysine-coated slides, which were dried at 37°C for 30 min and rehydrated for 10 min in PBS at room temperature (RT). For immunofluorescent labelling, the slides were incubated with blocking solution (10% normal goat serum, 1% bovine serum albumin in 0.3% PBS-Triton X 100) for 1 h at RT. The primary antibodies, anti-MCT2 (SLC16A7) (Alomone labs, Cat. No. AMT-012, RRID: AB_2340997), Anti-PKCα (Thermo Fisher Scientific, Cat. No. MA1-157, RRID: AB_2536865), and anti-calretinin (Abcam, Cat. No. A85366, RRID: AB_2748943) were diluted 1:100 in blocking solution and incubated at 4°C overnight. The slides were washed 3 times for 10 min each with PBS. Subsequently, the secondary antibody, diluted 1:350 in PBS, was applied to the slides, which were incubated for 1 h at RT. Finally, the slides were washed with PBS and covered in Vectashield with DAPI (Vector).

### TUNEL assay

Fixed slides from retinal explant cultures were dried at 37°C for 30 min and washed in phosphate-buffered saline (PBS) solution at room temperature (RT), for 15 min. Afterwards, the slides were placed in TRIS buffer with proteinase K at 37°C for 5 min to inactivate nucleases. The slides were then washed with TRIS buffer (10 mM TRIS-HCL, pH 7.4), 3 times for 5 minutes each. Subsequently, the slides were placed in ethanol-acetic acid mixture (70:30) at -20°C for 5 min followed by 3 washes in TRIS buffer and incubation in blocking solution (10% normal goat serum, 1% bovine serum albumin, 1% fish gelatin in 0.1% PBS-Triton X100) for 1h at RT. Lastly, the slides were placed in the terminal dUTP-nick-end labelling (TUNEL) solution (labelling with either fluorescein or tetra-methyl-rhodamine; Roche Diagnostics GmbH, Mannheim, Germany) in 37°C for 1 h and covered in Vectashield with DAPI (Vector, Burlingame, CA, USA) thereafter.

### Microscopy and cell counting

Fluorescence microscopy was performed with a Z1 Apotome microscope equipped with a Zeiss Axiocam digital camera (Zeiss, Oberkochen, Germany). Images were captured using Zen software (Zeiss) and the Z-stack function (14-bit depth, 2752*2208 pixels, pixel size = 0.227 µm, 9 Z-planes at 1 µm steps). The raw images were converted into maximum intensity projections using Zen software and saved as TIFF files. Inner retinal cells stained by the TUNEL assay were counted manually on 3 images per explant, the average cell number in each INL area was estimated based on DAPI staining and used to calculate the percentage of TUNEL positive cells.

### Retinal slice preparation for imaging and electrophysiological experiments

For FRET experiments, the explants were separated from the culture inserts and placed in a chamber with extracellular solution. For imaging and electrophysiology experiments, p30 mice were used. Eyes were enucleated and the retina was kept in extracellular solution during the remainder of the procedure. The eye was cut along the *ora serrata* to separate the anterior and posterior chambers, and the retina was separated carefully from the choroid – sclera. The extracellular solution for maintaining both types of slices contained (in mM): 119 NaCl, 23 NaHCO_3_, 1.25 NaH_2_PO_4_, 2.5 KCl, 2.5 CaCl_2_, 1.5 MgSO_4_, 20 glucose and 2 Na^+^ pyruvate, aerated with 95% O_2_ and 5% CO_2_, pH 7.4. Subsequently, the tissue was embedded in type VII agarose (Cat. No. 39346-81-1 Sigma Aldrich) dissolved in a solution composed of (in mM): 119 NaCl, 25 HEPES, 1.25 NaH_2_PO_4_, 2.5 KCl, 2.5 CaCl_2_, and 1.5 MgSO_4_ at a pH of 7.4. Finally, the tissue was cut with a vibratome (Leica VT1000S) to 200 μm thickness. The slices were transferred to the microscope recording chamber, sustained by a U-shaped platinum wire and superfused with oxygenated extracellular solution at room temperature (20°C) under photopic conditions.

### Calcium and sodium imaging

For imaging experiments, 200 µm retinal slices were obtained as previously described, and incubated for a period of 1 h in a dark room in 2 ml of extracellular solution containing 5 µM fluo- 4 AM (Thermo Fisher Scientific) or 7 µM Corona Green AM (Thermo Fisher Scientific) in 0.04% Pluronic Acid. Afterwards, the slices were imaged as previously described ^21^. Relative fluorescence intensities (ΔF/F0) of the regions of interest (ROIs) were obtained by dividing all images through the initial (pre-pulse) image of the series. In this study, all ROIs were chosen from the cell bodies of putative inner retinal neurons, located in the INL. To decrease the effect of photobleaching, periods of light stimulation lasted for 1 s and were followed by 10 s of darkness for the duration of the experiments. For MCT inhibition experiments, the retinal slices were incubated for 15 min with 0.1 µM SR-13800 (Tocris, Cat. No. 5431), 2 µM AR-C155858 (Tocris, Cat. No. 4960), 4 µM Shikonin (Sigma-Aldrich, Cat. No. S7576), or 15 µM FX-11 (MedChem Express, Cat. No. HY-16214) and then stimulated with 12 mM KCl through bath perfusion.

### Electrophysiology

Retinal slices from p30 *ex vivo* retinas were visualized with an upright microscope (Nikon Eclipse FN1) equipped with a 40x water-immersion objective, infrared differential interference contrast and a digital camera (TCH 1.4 LICE, Tucsen Photonics). Images were captured with ISCapture software and processed by Adobe Photoshop CS (Adobe Systems Incorporated). Patch clamp recordings were made from RBCs, whose tentative identity was corroborated by comparing the axon terminal stratification within the IPL after dialysis of Lucifer yellow through the patch pipette. Standard intracellular solution contained (in mM): 125 K^+^ gluconate, 10 KCl, 10 HEPES, 2 EGTA, 2 Na_2_ATP, 2 NaGTP and 1% Lucifer yellow. Recording electrodes were fabricated using borosilicate glass capillaries (1.5 mm OD, 0.84 mm ID; WPI) and pulled to resistances between 10 – 15 MΩ on a Flaming/Brown electrode puller (Sutter P-97). Experiments were only performed if the patch seal resistance was above 1 GΩ and series resistance below 30 MΩ. Signals were amplified with an EPC7- plus patch clamp amplifier (HEKA Elektronik), digitized and sampled at 10 kHz with an A/D board (Digidata 1550, Molecular Devices). Recordings were acquired using the software PClamp 10.4 (Molecular Devices).

For patch clamp experiments, the retinal slices were incubated for 15 min with either SR-13800 or AR-C155858 and then different protocols were applied. Between the experiments, cells were held at a resting membrane voltage of −60 mV. To measure outward currents, 10 mV voltage steps (200 ms duration) from −100 mV to 40 mV were applied to obtain the voltage-dependent current patterns. To measure voltage-activated Ca^2+^ currents, cells were patched with an internal solution containing (in mM): 90 Cs-methanesulfonate, 20 TEA-Cl, 10 HEPES, 10 EGTA, 10 Na_2_- phosphocreatine, 2 MgATP, and 0.2 NaGTP with pH adjusted to 7.4 with CsOH. To obtain the currents, cells were held at a resting membrane voltage of −60 mV and 5 mV voltage steps (100 ms duration) from −70 mV to +20 mV, were applied. Finally, to study the membrane potential, the current was clamped at 0 pA, and the average of 1 minute of recording was used to define the membrane potential.

### FRET measurements

At p11 and p12, the explants were transduced by overnight incubation with 5x10^6^ plaque-forming units (PFU) of Ad Laconic, AAV-Laconic or Δ6, and imaged after two weeks in cultures. Adenoviral serotype vectors encoding FRET nanosensor Ad FLII12Pglu-700Δ6 (Takanaga et al., 2008)) and Ad Laconic (San Martín et al., 2013) were a gift from Dr. Ivan Ruminot from the Centro de Estudios Científicos (CECs) in Valdivia, Chile. The AAV-GFAP-Laconic (Laconic: Addgene #44238; hGFAP promoter fragment: DOI: 10.1002/glia.20622) and AAV-hSYN-Laconic (Laconic: Addgene #44238) was constructed by the viral vector facility of ETH Zurich. Since the lactate and glucose sensors used here have similar excitation/emission spectra (Laconic: 460/492 nm for mTFP and 515/526 for Venus; Δ6: 440/480 nm for CFP and 513/530 nm for YFP), retinal slices were excited at 430 nm and visualized at 480 nm and 530 nm peak wavelength, as previously reported (San Martín et al., 2013; Barros et al., 2014). All experiments were performed at room temperature (22– 25°C) with an upright fluorescence microscope (Olympus BX51) equipped with a 40x water- immersion objective, an Optosplit II emission image splitter (Cairn, UK), and a Sensicam QE digital camera (Cooke Corp.).

FRET experiments were performed in extracellular solution containing (in mM): 119 NaCl, 23 NaHCO_3_, 1.25 NaH2PO_4_, 2.5 KCl, 2.5 CaCl_2_, 1.5 MgSO_4_, 5 glucose,1 lactate and, aerated with 95% O_2_ and 5% CO_2_, pH 7.4. The lactate and glucose concentrations were chosen not to saturate the FRET sensors. Data acquisition was performed by custom software written in Python 4.0.1. At the end of the experiments, data were exported for off-line analysis of fluorescence intensities from each channel. To obtain the FRET ratio for Laconic (mTFP/Venus) and Δ6 (YFP/CFP), fluorescence intensity values from each ROI and background were measured in ImageJ, version 1.52p (NIH, RRID: SCR_003070). In this study, all ROIs were chosen from the somas of MCs and putative inner retinal neurons. The FRET data are displayed as the relative FRET ratio, in percentage of change over time of single experiments.

To calculate the delta ratio between the depletion and the saturation for both sensors, the minimum response (depletion) was obtained during the application of 10 mM pyruvate, and the maximum response (saturation) during 10 mM lactate stimulation, their difference yielding the delta ratio. To evaluate the effect of different MCT inhibitors on lactate influx, the positive amplitude peak was measured before and after drug application. The decay time of these responses was calculated as the time required to return to baseline from the response peak. In both cases, the lactate response and the decay time were first normalized per cell based on the control response and then averaged throughout the experiments. Finally, the slopes of the responses were calculated by fitting a linear regression and obtaining the slope via the lineal function: *y= mx + b*, where ***m*** is the slope. Statistical analysis was performed using GraphPad Prism software (RRID:SCR_002798).

### Pharmacology

Since there is only a limited number of commercial MCT inhibitors available (Murray et al., 2005; Ovens et al., 2010; Vélez et al., 2021), we used three potent and specific inhibitors to isolate MCT isoforms: SR-13800 for MCT1 (SR; Tocris, Cat. No. 5431), AR-C155858 (AR-C; Tocris, Cat. No. 4960) for MCT1 and MCT2, and Syrosingopine (Syro; Sigma Aldrich, Cat. No. SML1908) to inhibit MCT1 and MCT4. To study intracellular enzymes involved in lactate production, we used Shikonin (Sigma Aldrich, Cat. No. S7576) to inhibit the M2 isoform of pyruvate kinase (PK), and FX-11 to inhibit lactate dehydrogenase A (Medchem Express, Cat. No. HY-16214).

### Data analysis

All data were first analyzed for normality using the Shapiro–Wilk test. If the test determined that the data did not conform to a normal distribution, significant differences were established with the Wilcoxon Matched-Pairs Signed Ranks Test, and Kruskal-Wallis test followed by a Dunńs multiple comparison test. Whether the Shapiro-Wilk test determined that the data conformed to a normal distribution, significant differences were established with the paired t-test and one-way ANOVA multiple comparison test, followed by Tukey’s multiple comparisons test. The α value was set to 0.05. Unless otherwise stated, data values are given as mean ± SD. Significance levels as indicated by asterisks were: * p < 0.05; **p < 0.01, *** p < 0.001. Statistical analysis was performed using GraphPad Prism software (RRID:SCR_002798).

## Supplementary figures

**Figure S1.**
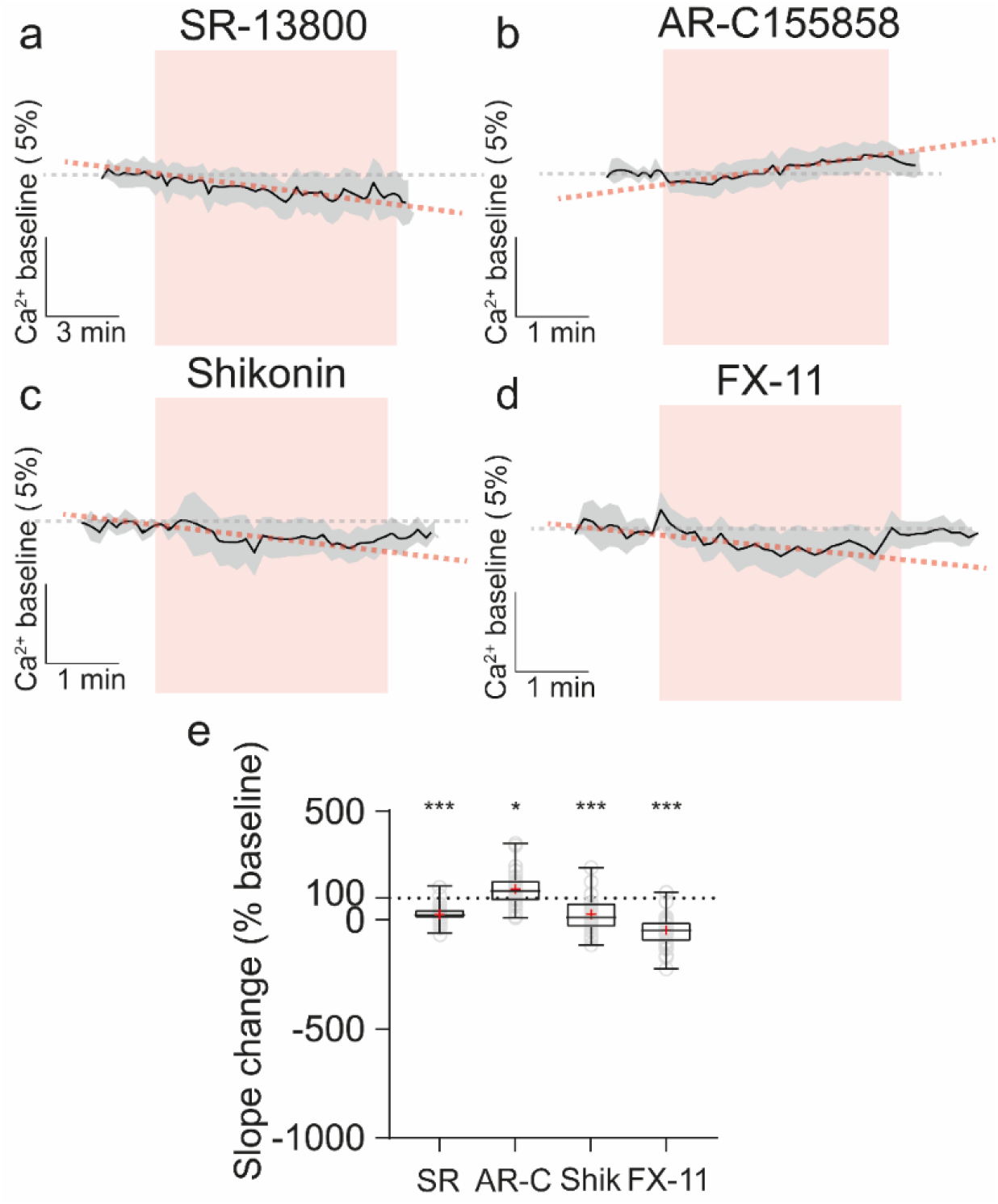
Alterations in relative basal calcium levels during impaired lactate metabolism. **(a-d)**. Traces of an experiment in each condition (black line=mean, gray shade=standard deviation, number of cells per experiment: SR = 10 cells, AR-C = 19 cells, shikonin = 15 cells, FX- 11 = 22 cells), indicating the incubation with different drugs (red boxes). The gray dashed line shows the slope in the pre-incubation condition, while the red dashed line represents the change in slope during drug incubation. **(e)** Statistical analysis of basal calcium levels. Box plot showing significant alterations in all conditions: Under inhibition of MCT1 (SR; p < 0.0001), PKM-2 (Shik; p < 0.0001) and LDH-A (FX-11; p < 0.0001), we observed a significant reduction of basal calcium levels. However, inhibition of MCT2 (AR-C; p = 0.0148) revealed an increase in relative calcium levels. Control is represented as dashed line at 100%. Graphs show median ± minimum and maximum values and mean in red. Individual values are shown as open circles (gray). Asterisks indicate * p < .05, ***p < .001. SR = MCT1 inhibitor; AR-C = MCT2 inhibitor; Shikonin = PKM2 inhibitor; FX-11 = LDH-A inhibitor.

**Figure S2.**
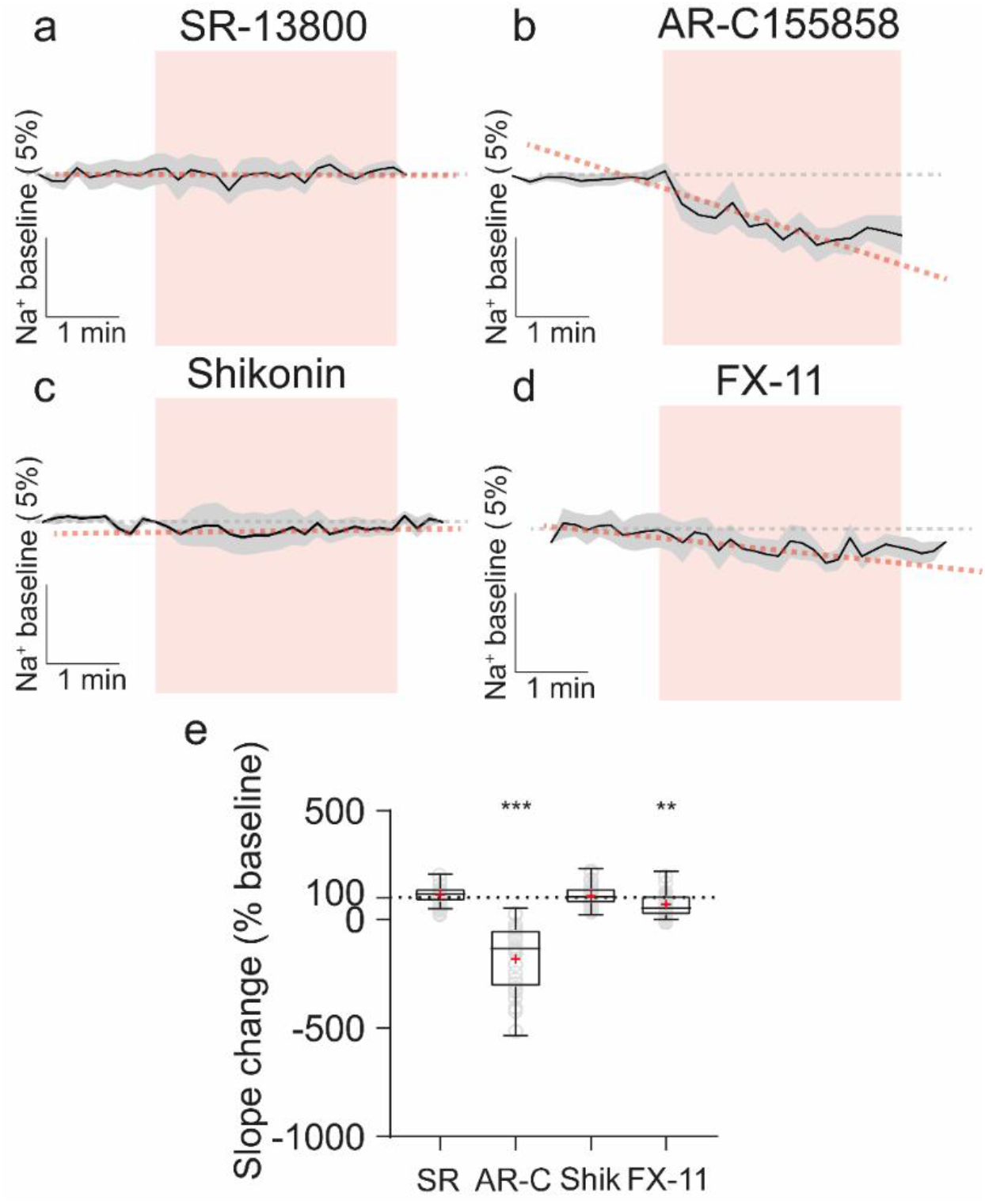
Disruption in lactate metabolism impaired basal sodium levels. **(a-d).** Traces of an experiment in each condition (black line = mean, light gray shadow = standard deviation, number of cells per experiment: SR = 12 cells, AR-C = 13 cells, Shikonin = 12 cells, FX-11 = 17 cells), indicating with incubation under different drugs (red boxes). The gray dashed line shows the slope in the pre-incubation condition, while the red dashed line represents the change in slope during incubation with drugs. **(e)** Statistical analysis of basal sodium levels. Box plots showed alterations only under two conditions: Under the inhibition of MCT2 (AR-C; P <0.0001) and slightly under LDH-A inhibition (FX-11; P = 0.0014). While under the blocking of MCT1 (SR; P = 0.0602) and PKM2 inhibition (Shik; P = 0.19) the basal sodium level was not affected. Control is represented as dashed line at 100%. Graphs show median ± minimum and maximum values and mean in red. Individual values are shown as open circles (gray). Asterisks indicate * p < .05, **p< .01. SR= MCT1 inhibitor; AR-C = MCT2 inhibitor; Shikonin = PKM2 inhibitor; FX-11 = LDH-A inhibitor.

**Fig. S3.**
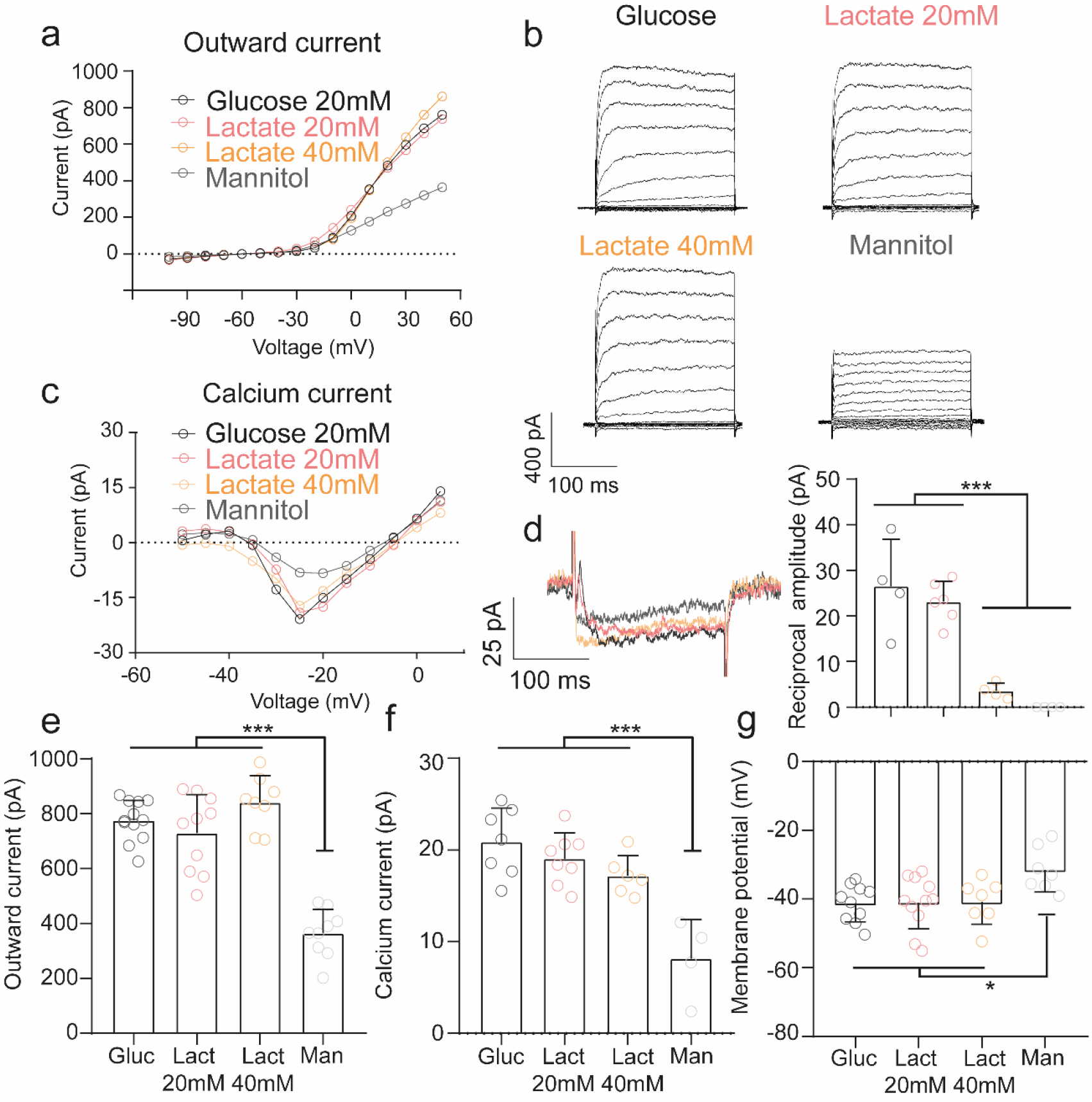
RBCs can use lactate in the absence of glucose to maintain their current profiles. **(a, c)**, Comparison of the current-voltage relationship of the outward currents and calcium currents under different pharmacological conditions. **(b, d),** Representative responses to depolarizing voltage steps. The reciprocal feedback current was altered only in the mannitol (p = 0.0002), lactate 40 mM condition (p < 0.0001), but was unaffected in lactate 20 mM (p = 0.7856). **(e, f)**, In the absence of glucose, only in the mannitol condition a decrease in the outward currents and calcium currents (p < 0.0001) was observed (p < 0.0001), but no differences were noted in the lactate 20 mM and 40 mM conditions either in the outward currents (p = 0.7079; p = 0.4973) or calcium currents (p = 0.6924; p = 0.2019). **(g)**, Similar results were obtained when we measured the membrane potential, which displayed a depolarization in the mannitol condition (p = 0.0256), while it remained unaltered in lactate 20 mM and 40 mM (p = 0.9999; p = 0.9993). The data were analyzed by ordinary one-way ANOVA, with a subsequent Tukey’s multiple comparison post hoc test. Each dot reflects a single recorded cell. Graphs display the mean ± SD; asterisks indicate * p < .05, **p < .01, ***p < .001.

**Fig. S4.**
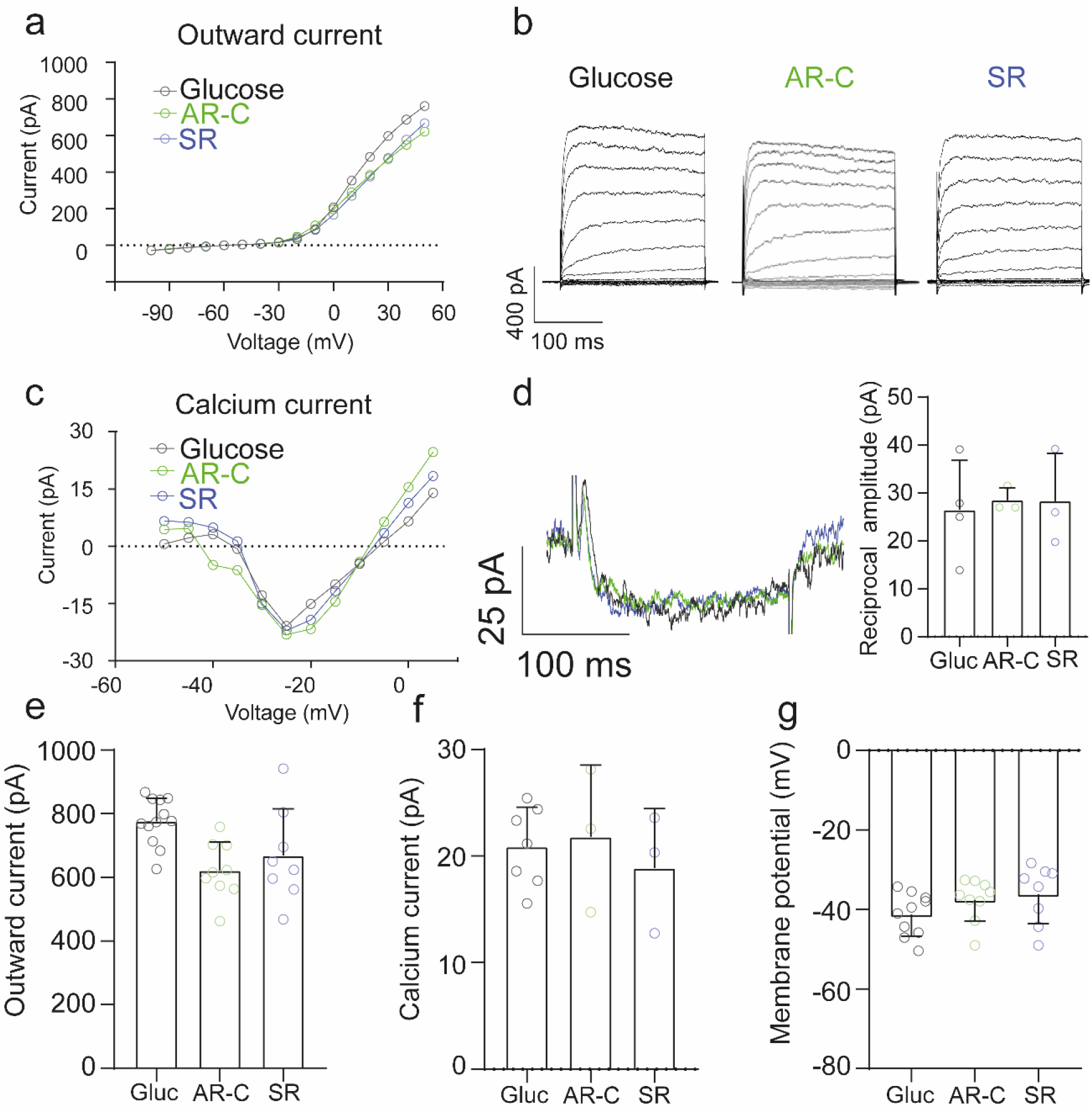
The inhibition of lactate transport does not affect RBC voltage-gated currents in presence of glucose. **(a, c)**, Comparison of voltage-current relationship of the outward currents and calcium currents in the different conditions. **(b, d),** Representative responses to depolarizing voltage steps. In the presence of glucose, the reciprocal feedback current was not altered in the AR-C (p = 0.9501) and SR condition (p = 0.9592). **(e, f)**, Likewise, neither AR-C (p = 0.0913) nor SR (p = 0.0693) affected the outward and calcium currents (p = 0.9581 and p = 0.8267, respectively). **(g)**, Similar results were obtained regarding the membrane potential, with no change caused by either AR-C (p = 0.3938) or SR (p = 0.1921). The data were analyzed by one-way ANOVA, with Tukey’s multiple comparison post hoc test. Each dot reflects a single recorded cell. Graphs display the mean ± SD. SR = MCT1 inhibitor; AR-C = MCT1 and MCT2 inhibitor

**Fig. S5.**
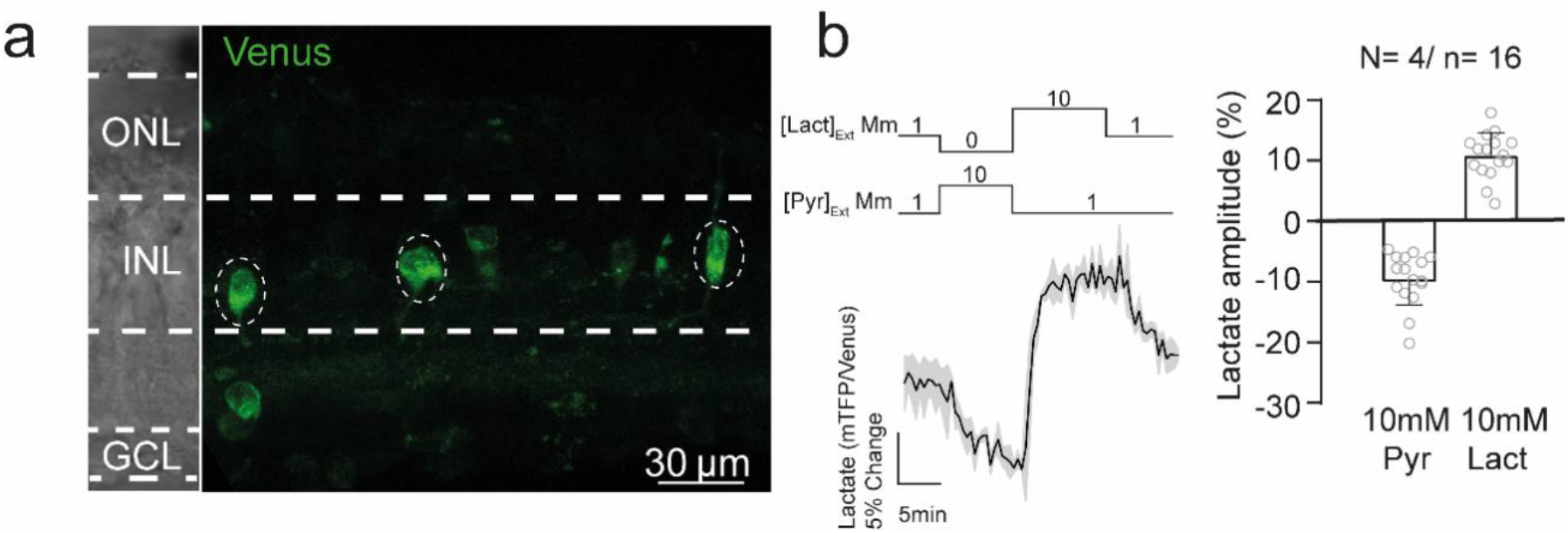
The Laconic nanosensor can be functionally expressed in inner retinal neurons. **(a)** Confocal image of Laconic expression in retinal explants after two weeks in culture. Dashed circles show the recorded area. **(b)** Dynamic range of the lactate sensor in inner retinal neurons. ONL= outer nuclear layer; INL = inner nuclear layer; GCL = ganglion cell layer. The black trace represents the average of one experiment (3 cells recorded), while the light grey shadow represents the standard deviation. The number of experiments is represented as: N = number of explants; n = number of cells recorded. Graph displays the mean ± SD.

**Figure S6.**
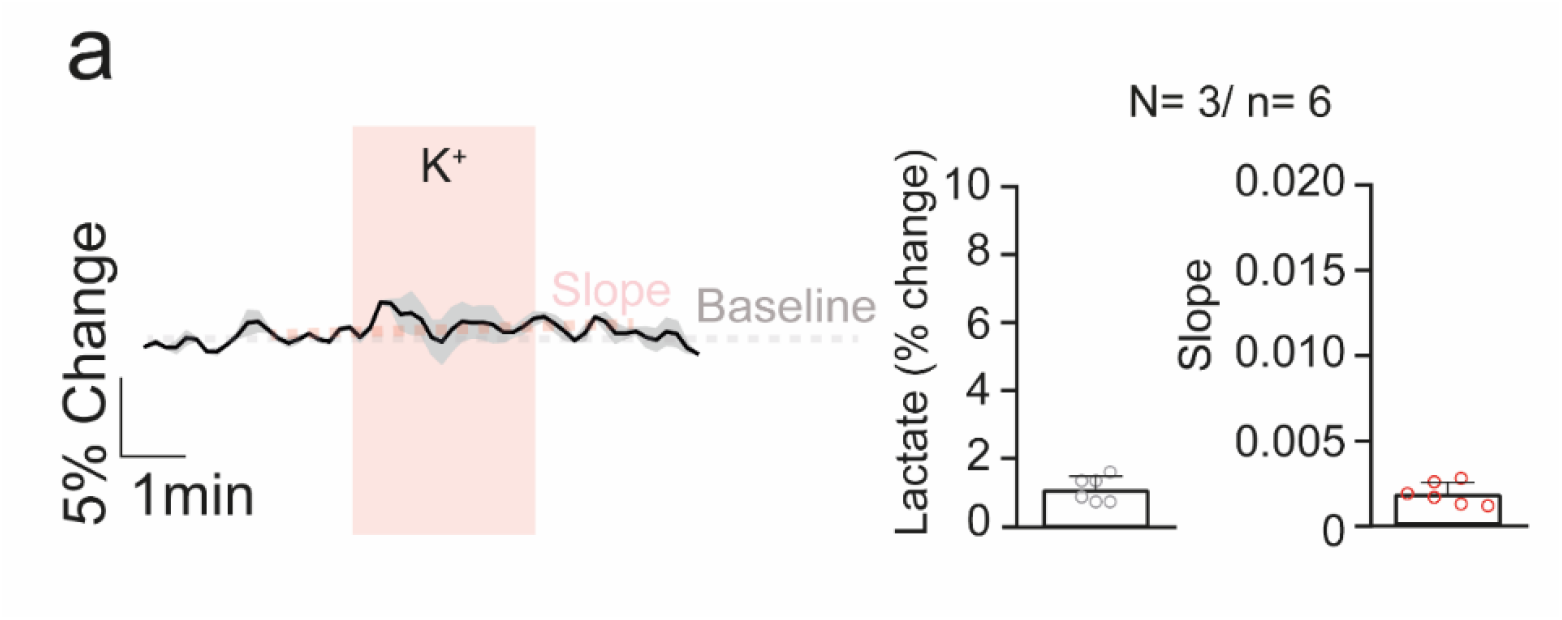
Depolarization by bath application of KCl has no effect on a group of inner retinal cells. **(a, left)** Representative traces of one experiment showing no responses during and after depolarization. The black trace represents the average of one experiment, while the light grey shadow represents the standard deviation. **(a, right)** Quantification of both amplitude and slope during the bath application of potassium does not demonstrate any alteration of these parameters. Graphs display the mean ± SD. The number of experiments is represented as: N = number of explants; n = number of cells recorded.

## Notes

### Competing Interest Statement

The authors have declared no competing interest.

